# Integration of Structured Biological Data Sources using Biological Expression Language

**DOI:** 10.1101/631812

**Authors:** Charles Tapley Hoyt, Daniel Domingo-Fernández, Sarah Mubeen, Josep Marin Llaó, Andrej Konotopez, Christian Ebeling, Colin Birkenbihl, Özlem Muslu, Bradley English, Simon Müller, Mauricio Pio de Lacerda, Mehdi Ali, Scott Colby, Dénes Türei, Nicolàs Palacio-Escat, Martin Hofmann-Apitius

## Abstract

**Background:** The integration of heterogeneous, multiscale, and multimodal knowledge and data has become a common prerequisite for joint analysis to unravel the mechanisms and aetiologies of complex diseases. Because of its unique ability to capture this variety, Biological Expression Language (BEL) is well suited to be further used as a platform for semantic integration and harmonization in networks and systems biology.

**Results:** We have developed numerous independent packages capable of downloading, structuring, and serializing various biological data sources to BEL. Each Bio2BEL package is implemented in the Python programming language and distributed through GitHub (https://github.com/bio2bel) and PyPI.

**Conclusions:** The philosophy of Bio2BEL encourages reproducibility, accessibility, and democratization of biological databases. We present several applications of Bio2BEL packages including their ability to support the curation of pathway mappings, integration of pathway databases, and machine learning applications.

**Tweet:** A suite of independent Python packages for downloading, parsing, warehousing, and converting multi-modal and multi-scale biological databases to Biological Expression Language

## 1. Background

The integration of heterogeneous, multi-scale, and multi-modal biomedical data has become a cornerstone of modern computational investigation of the mechanisms and aetiologies underlying complex diseases (Iyappan *et al.*, 2014; van Dam *et al.* 2014; Wanichthanarak *et al.*, 2015; Himmelstein et al., 2017; Fan *et al.*, 2019). An overarching strategy was proposed by Davidson *et al.* more than two decades ago that outlined the transformation of data into a common model, semantic alignment of related objects, integration of schemata, and federation of data (Davidson *et al.*, 1995). However, integration remains a challenging task that requires the identification and deep understanding of biological data sources and their respective formats, conversion, harmonization, and unification.

Initial interest in the semantic web and linked open data along with the adoption of RDF (Resource Description Framework^1^) in the biomedical community led to the Bio2RDF project, in which pipelines for converting and serializing several biological data sources to RDF were developed (Belleau *et al.*, 2008). Several updates have been issued since its deployment such as the inclusion of chemical information systems (Chen *et al.*, 2010). Further, it has also influenced in and has been adopted by subsequent projects such as Open PHACTS (Williams *et al.*, 2012). While RDF is highly expressive and each of these projects have developed and enforced well-defined schemata, the format is often not well-suited for downstream analyses and must first be queried with languages like SPARQL (SPARQL Query Language for RDF^2^) and subsequently be transformed into appropriate formats with general-purpose programming languages. Alternatives to RDF/SPARQL such as property graphs (e.g., Neo4j^3^, OrientDB^4^) are comparable (Alocci *et al.*, 2015) but also necessitate similar post-processing.

Conversely, there have been several biologically meaningful integration efforts (e.g., STRING; Warde-Farley, *et al.* 2010, GeneMANIA; Szklarczyk *et al.*, 2015, GeneCards; Stelzer *et al.*, 2016). However, most suffer from a lack of defined schemata or standardized data format that impede biological database interoperability. As interoperability itself is a multifaceted concept, we would like to highlight three of its facets: first, data sources should refer to named entities using high-quality, publicly accessible terminologies as prescribed by the Minimal Information Requested in the Annotation of Biochemical Models standard (Laibe and Le Novère, 2007). Second, data sources should additionally denote the ontological classes of named entities (e.g., gene, transcript, protein, pathway, disease) along with their reference using controlled vocabularies such as the Systems Biology Ontology (Courtot *et al.*, 2011). Some identifiers, such as those for genes, are often used to refer not only to the physical region of DNA within the genome, but also the corresponding RNA transcript(s) or protein product(s). Unfortunately, many biological databases do not explicitly distinguish between these entity classes. For example, the STRING database lists gene-centric homology relationships, transcript-centric co-expression relationships, and protein-centric protein-protein interactions using gene-centric nomenclature. While it may be possible to identify the classes of entities based on their incident relationships, doing so requires specific knowledge of the database including the semantics of its relationships. Third, resources should, at a minimum, map their relationships to controlled vocabularies such as the Relation Ontology^5^, or further use standardized data formats with defined semantics (e.g., PSI MI-TAB^6^) to minimize both the interpretation and implementation effort when combining them with other resources.

OmniPath (Türei *et al*., 2016) began to address these facets when it combined several signaling pathway and transcriptional regulation databases. It achieved interoperability between several databases by normalizing the identifiers and relationships between entities from several databases describing the same phenomena (e.g., microRNA-target interactions, protein-protein interactions, etc.) and creating a unified network. However, because it did not use a standard format or schema as mentioned in the third facet for interoperability, OmniPath itself cannot readily be directly integrated with other biological data sources. Pathway Commons (Cerami *et al.*, 2011) addressed this concern when combining several molecular pathway and interaction databases by translating the source databases into the BioPAX standard (Demir *et al.*, 2010) using automated pipelines. However, it suffers from low granularity and low recovery of information from some of its primary biological data sources which may be due to prioritization of software development time, data usage restrictions, or shortcomings in the BioPAX standard. While BioPAX is well-suited for representing biological reactions and transformations, it is limited in its ability to represent correlative and associative relationships across multi-scale biology (e.g., at the levels of processes, phenotypes, and clinical observations).

As an alternative, we propose the use of Biological Expression Language (BEL; Slater, 2014) as an integration schema in order to overcome the limits faced by previous efforts and to simultaneously address all three facets of interoperability. BEL has begun to prove itself as a robust format in the curation and integration of previously isolated biological data sources of high granular information on genetic variation (Naz *et al.*, 2016), epigenetics (Irin *et al.*, 2015), chemogenomics (Emon *et al.*, 2017), and clinical biomarkers (Iyappan *et al.*, 2017). Its syntax and semantics are also appropriate for representing, for example, disease-disease similarities, disease-protein associations, chemical space networks, genome-wide association studies, and phenome-wide association studies.

With the same focus on reproducibility as Bio2RDF, OmniPath, and Pathway Commons as well as deference to software maintainability and the ease of development and inclusion of new biological data sources, we have developed a growing list of *Bio2BEL* packages, each capable of downloading, structuring, and serializing various biological data sources to BEL (**Table 2**). Each can be found in the Bio2BEL GitHub organization (https://github.com/bio2bel) as an independent open-source Python package that can readily be installed with pip. We have also developed and freely provided a framework (https://github.com/bio2bel/bio2bel) in the Python programming language to enable code reuse and the fast generation of additional Bio2BEL packages. Notably, the list of Bio2BEL packages includes one for OmniPath as a proof of concept that authors of other resources can implement their own Bio2BEL packages. In this article, we present the philosophy and implementation of Bio2BEL packages, a summary of past and future Bio2BEL packages, and finally, several case studies including the utility of Bio2BEL packages during curation of pathway mappings, in the analysis of cancer genome data, and for machine learning applications.

## 2. Implementation

Bio2BEL comprises numerous independent open-source Python packages that each enable reproducible access to a given biological data source (**Figure 1**). Each Bio2BEL package contains five components: 1) a definition of the source database or knowledge base, 2) an automated downloader for the data, 3) a parser for the data, 4) a storage and querying system for the data, and 5) a protocol for serializing the data to BEL (**Figure 2**). In this section, we outline the components of a Bio2BEL package and their implementation details.

**Figure 1:**
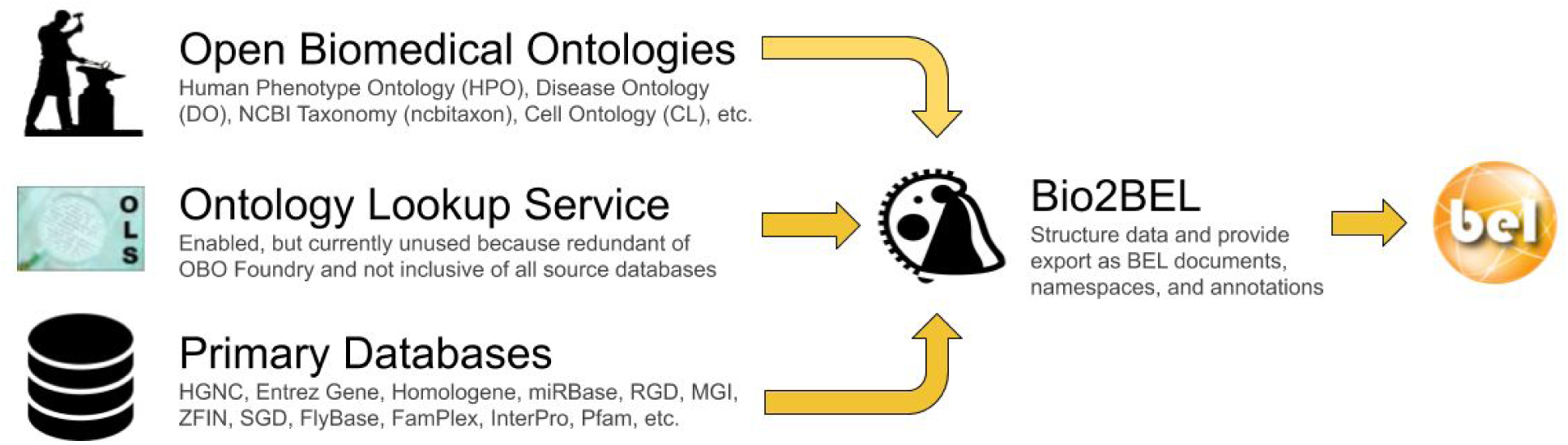
Though their main focus is on generating BEL documents, some Bio2BEL repositories have secondary goals of generating the BEL namespace and annotation files necessary to support manual curation. Most rely on primary databases, but the Bio2BEL framework also includes functions for generating them from standard Open Biomedical Ontology documents, or through the EBI Ontology Lookup Service (Cote *et al*., 2006). Logos adapted from http://obofoundry.org, https://www.ebi.ac.uk/ols, and https://openbel.org.

**Figure 2:**
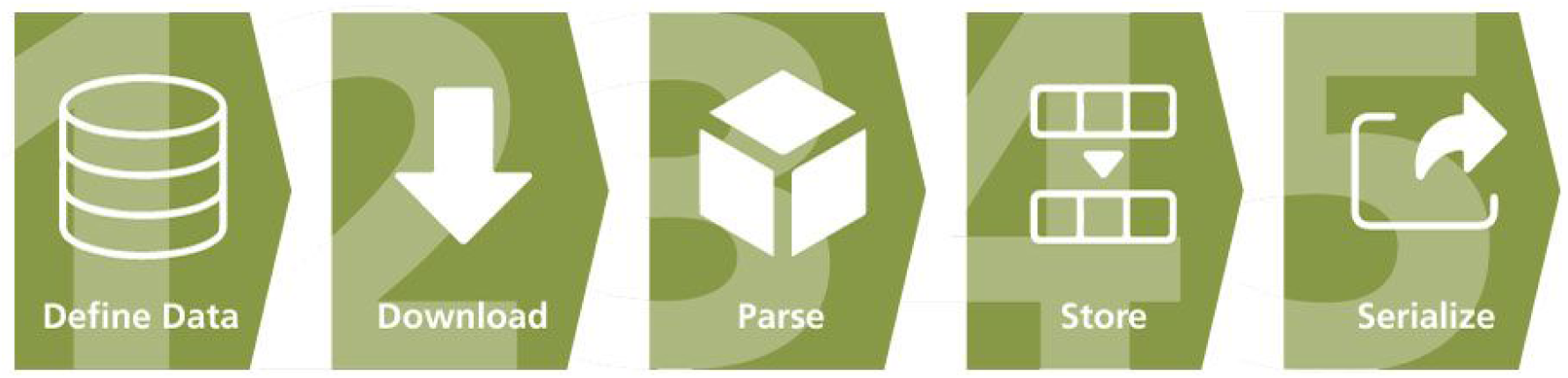
A graphical overview of the sequentially ordered components of a Bio2BEL package. These components correspond to the philosophy that reproducibility and accessibility can ultimately lead to the democratization of the usage of prior biological knowledge.

### 2.1. Components of a Bio2BEL Package

As this section outlines the core components and philosophy of a Bio2BEL package, it illustrates the tasks and thought process of a scientific software developer as they implement a new Bio2BEL package.

#### 1. Definition of Data

The first step in generating a Bio2BEL package is to understand the source data. This requires determining if the data are publicly accessible, if they are versioned (and how the location changes with versions), and if they are available under a permissive license. Bio2BEL packages do not contain data themselves and only refer to the locations of the original data sources. For those that are versioned, providers commonly generate symlinks to the most recent version (e.g., InterPro; ftp://ftp.ebi.ac.uk/pub/databases/interpro). These characteristics help minimize licensing issues while enabling the resulting packages to update their content without changing code. Then, the developer implements custom code that makes the appropriate interpretations to convert the source data to BEL. Below, three types of data that can be readily integrated in BEL are described along with accompanying **Table 1**.

**Table 1.**
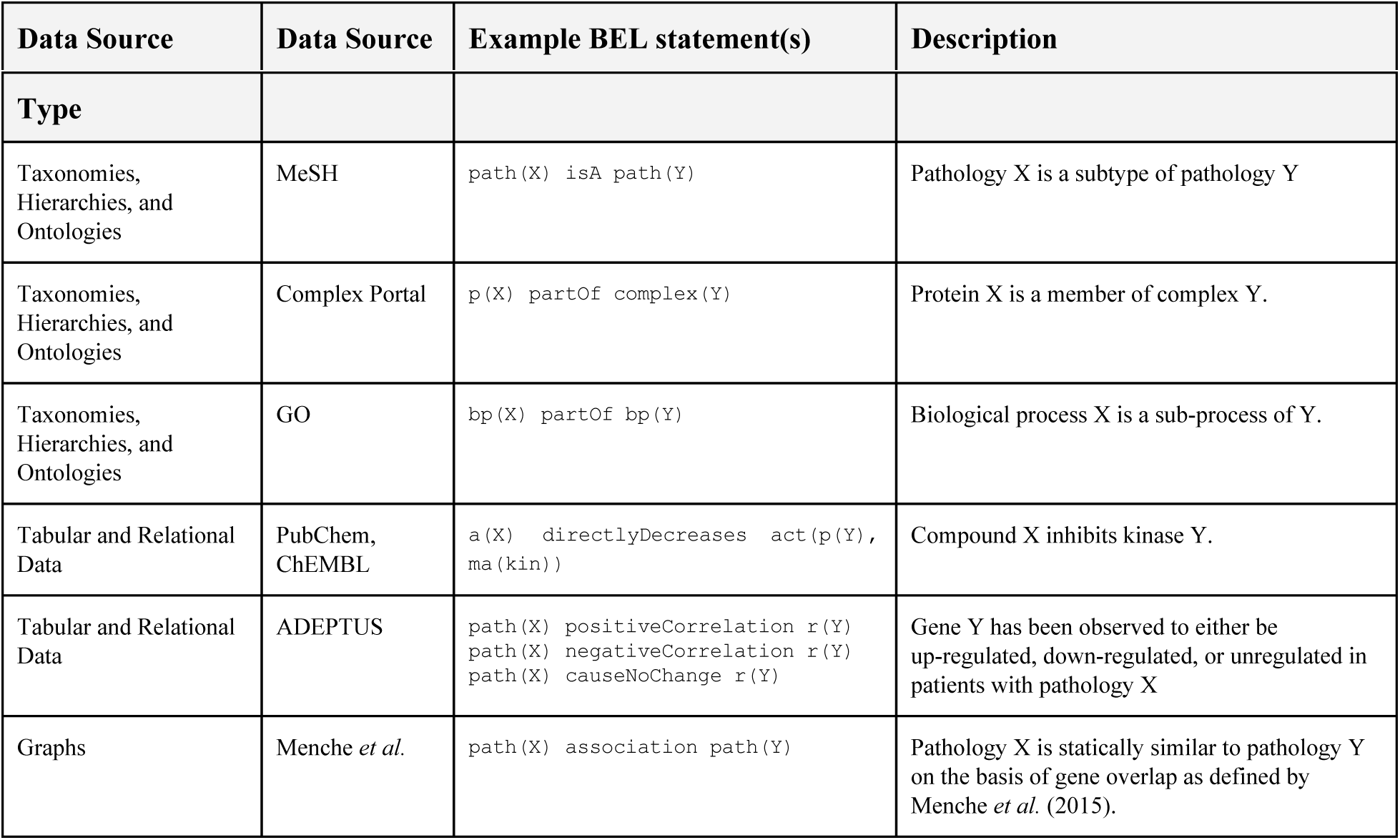
Example BEL statements generated by several different types of data sources

##### I. Taxonomies, Hierarchies, and Ontologies

The Medical Subject Headings (MeSH; Rogers, 1963) multi-hierarchy can be converted to BEL by generating an *isA* relationship between each MeSH descriptor and all of its corresponding parents in the associated MeSH tree. Nomenclatures like the Complex Portal (Meldal *et al*., (2015) also define *partOf* relations between protein complexes and their substituents. The multi-hierarchy in Gene Ontology (GO; Carbon *et al.*, 2017) can be converted similarly, which contains both *isA* relations and *partOf* relations.

##### II. Tabular and Relational Data

Enzyme inhibitors from ChEMBL and PubChem can be encoded like *a(X) directlyDecreases act(p(Y), ma(kin))*, and disease-specific differential gene expression can be encoded like *path(X) positiveCorrelation r(Y)* or *path(X) negativeCorrelation r(Y)*, or *path(X) causeNoChange r(Y)* depending on whether the gene’s expression is up-regulated, down-regulated, or not regulated, respectively. Further, BEL relationships can be extended include metadata (i.e., annotations) describing their quantitative aspects. For example, IC_50_, EC_50_, or other kinetic assay measurements as well as provenance and biological contextual information (e.g., original publication, cell line, assay type) can be included with the enzyme inhibition relationships from ChEMBL. Similarly, the log_2_ fold change and *p*-values can be included with relationships about differential gene expression.

##### III. Graphs

Wet-laboratory experimentation can be used to generate networks of directly observed phenomena (e.g., protein-protein interaction networks) and indirectly observed phenomena (e.g., gene co-expression networks). Graphs are often distributed as tabular data to include additional information about their constituent nodes and edges and there is often overlap with the previous data type describing tabular and relational data. *In silico* experimentation can also be used to derive edges from experimental data sets or even other graphs. For instance, bipartite graphs can be projected to homogeneous graphs consisting of a single entity and edge type as suggested by Sun *et al.* (2014). Menche *et al.* (2015) used this strategy and computed a homogenous graph of disease-disease associations from a bipartite graph of diseases and their associated genes.

#### 2. Downloader

The Bio2BEL framework follows a functional programming paradigm to provide an abstraction of the acquisition of data over common internet protocols like HTTP, HTTPS, and FTP. With only the URL of the data set as an input, Bio2BEL generates a download function that wraps Python’s built-in *urllib* module and a simple caching mechanism in the local filesystem that avoids unnecessary network usage and duplication of potentially large files. However, some data sources, such as DrugBank (Wishart *et al.*, 2018), are not available without authentication and cannot make use of this abstraction. In those cases, developers can substitute the standard code provided in the Bio2BEL framework with custom implementations. We have taken this route for several of the packages presented in the Results section of this paper for repositories including DrugBank and MSigDB (Liberzon *et al.*, 2015).

#### 3. Parser

There are several common file formats used by biological data sources (e.g., CSV, TSV, XML, RDF, JSON, KGML^7^, Stockholm^8^, OBO^9^, OWL^10^). Data may also (and sometimes only) be accessible through public application programming interfaces (APIs) such as the data from KEGG (Kanehisa *et al.*, 2017), Reactome (Fabregat *et al.*, 2018), and BioThings (Xin *et al*., 2016). Alternatively, data may be available through software packages usage such the Affymetrix R package (Gautier *et al.*, 2004) and HaploReg (Ward and Kellis, 2012). After each Bio2BEL package’s downloader generates a local copy of the data, the developer can either use one of the pre-defined parser functions from the Bio2BEL framework or implement a custom parser. For the most simple formats (i.e., CSV and TSV), the Bio2BEL framework automatically generates a parser that uses the *pandas* package (McKinney, 2010; https://github.com/pandas-dev/pandas). Formats like XML, JSON, and Stockholm have corresponding parsers built into the Python language or standard biology-focused packages, but the information contained within often needs custom logic for restructuring such as in the case of KGML, BioPAX, or PSI MI-XML ^11^. The remaining custom formats all require custom parsers and logic. We have already implemented Bio2BEL that used CSV and TSV data (e.g., InterPro, ExCAPE-DB), XML (e.g., DrugBank), RDF (e.g., WikiPathways), JSON and KGML (e.g., KEGG), Stockholm (e.g., miRBase), and OBO and OWL (e.g., GO, DOID).

In the case of tabular data, the developer has the opportunity to annotate the column headers and their corresponding data types, which are not always included in the data and may be sought from various readme files or by exploring the corresponding website. Further, the contained data might be more useful after normalization or augmentation with information from other biological data sources. Because some databases provide identifiers with redundant information, such as the duplication of the namespace in the identifier, they must be normalized. For example, each identifier in the Disease Ontology (Schriml *et al*., 2018) is prefixed by its namespace, DOID, as can be seen in the Compact URI for the entry for restless legs syndrome, DOID:DOID:0050425. In the corresponding Bio2BEL DOID package, as well as those for others (e.g., HGNC, Gene Ontology) we normalized these identifiers to remove the redundant information. Because the main Entrez Gene database does not contain crucial information for genes, such as their chromosomal coordinates in various genomic builds, we augmented the data in the Bio2BEL Entrez package for each gene with information from RefSeq so that the genomic positions and corresponding genome build for each gene were readily accessible. Additionally, several databases that reference genes only use their HGNC gene symbols and not stable identifiers, and therefore require this additional normalization step.

#### 4. Storage

Though this step may be considered optional after parsing the data, it is helpful for future reuse to choose a database type and develop a schema with which the data can be stored. Often, relational databases that can be queried with SQL are an appropriate choice. The Bio2BEL framework provides a full harness for generating an object-relational mapping (ORM) using the SQLAlchemy (https://www.sqlalchemy.org) Python package that handles generation of the SQL schema and storage of the data in a SQL database. Corresponding entity-relation diagrams can be found in the supplementary data repository at https://github.com/bio2bel/bio2bel-manuscript-supplement. While all Bio2BEL packages have, until now, used SQL databases with the SQLAlchemy ORM, there exists alternatives such as graph databases built on RDF or property graphs like Neo4J or OrientDB with a corresponding object-graph mapper that have been successfully employed in downstream applications using biological knowledge graphs (Himmelstein *et al.*, 2017; Saqi *et al.*, 2018).

#### 5. Serializer

The final aspect of a Bio2BEL package is either to serialize the parsed data as BEL or to export the accompanying database as BEL. Entities in the SQL database that correspond to nodes and edges in BEL graphs can be converted by extending their respective ORM classes with Python functions using the internal domain-specific language provided by PyBEL (Hoyt *et al.*, 2018a). It can then be output in several formats provided by PyBEL and its growing ecosystem of plugins as well as it shields Bio2BEL packages from changes to the BEL language. Additionally, some Bio2BEL packages wrap standard nomenclature resources such as HGNC (Yates *et al*., 2017) and are able to generate BEL namespace files that are a necessary in both manual and automated curation of content in BEL (**Figure 2**). This step is deeply connected with the prior step related to the definition of the data.

### 2.2. Implementation Details

The Bio2BEL framework and Bio2BEL packages are implemented in Python with accessibility and readability in mind. The framework provides an abstract class *bio2bel.Manager* whose functionality all Bio2BEL packages must completely implement. Using these definitions, the framework automatically generates a uniform command line interface (CLI) that includes functions for populating the database, clearing the database, reloading data from the source, generating a web application with a view over the contents of the database, and serializing to BEL.

The Bio2BEL framework and Bio2BEL packages use flake8 (https://github.com/PyCQA/flake8) to enforce code quality, a setup.cfg file to describe the package, setuptools (https://github.com/pypa/setuptools) to build distributions, pyroma (https://github.com/regebro/pyroma) to enforce package metadata standards, sphinx (https://github.com/sphinx-doc/sphinx) to build documentation, Read the Docs (https://readthedocs.org) to host documentation, pytest (https://github.com/pytest-dev/pytest) as a testing framework, coverage (https://github.com/nedbat/coveragepy) and Codecov (https://codecov.io) to monitor testing coverage, and Travis-CI (https://travis-ci.com) as a continuous integration service. Further, we provide a template for Cookiecutter (https://github.com/audreyr/cookiecutter) at https://github.com/bio2bel/bio2bel-cookiecutter such that the structure of new packages can be quickly generated containing all of the configuration for each of these tools.

### 2.3. Implications of the Bio2BEL Philosophy

Because all Bio2BEL packages are uniform in their implementation and CLI usage, it is trivial to provide a Dockerfile and Docker-Compose configuration for quick deployments. In the future, we plan to automatically generate RESTful APIs, which may be more useful to deploy internally than to use publicly available ones due to constraints like rate-limits. Because all Bio2BEL packages are independent, they avoid two major problems of monolithic codebases: they are more robust to breakages or failures in a single package and they can be installed as needed, which is pertinent as the data sources become larger, more heterogeneous, and more complex.

Further, Bio2BEL packages can be generated by any group, and registered with the Bio2BEL framework using Python entry points (https://packaging.python.org/specifications/entry-points) that can be defined in the installation configuration. While the Cookiecutter template allows new developers to quickly generate a package with the correct format, a full tutorial for implementing a uniform Bio2BEL package can be found at https://bio2bel.readthedocs.io/en/latest/tutorial.html.

## 3. Results

After describing the Bio2BEL framework and the requirements for implementing new Bio2BEL packages, we present a list of the independent Bio2BEL packages that we have already implemented in **Table 2**. We note that several of the data sources have already been included in other meta-databases like Pathway Commons and Bio2RDF, but we have chosen to implement the Bio2BEL packages using the source data rather than deriving results from these databases to provide a complementary resource for those familiar with and interested in using BEL. This choice also reduces dependencies on other projects that may not be maintained and protects against data loss during multiple conversions.

**Table 2.** A non-exhaustive list of biological data sources already available as Bio2BEL packages

| Name | Description | Terms | Relations |
| --- | --- | --- | --- |
| adeptus | Disease-specific differential gene expression |  | 4,943 |
| chebi | Chemical multi-hierarchy | 138,863 |  |
| compath | Pathway-pathway equivalences and hierarchies |  | 1,795 |
| ddr | Disease-disease relationships |  | 2,997 |
| drugbank | Drug-target interactions | 11,292 | 25,199 |
| entrez | Genes and orthologies | 388,986 |  |
| excape | Chemical-target interactions |  | 70,850,163 |
| expasy | Enzyme classification and membership | 6,718 | 243,914 |
| famplex | Protein family and complex hierarchy |  | 4,462 |
| flybase | Drosophila gene nomenclature and orthologies | 245,565 |  |
| go | Biological process multi-hierarchy | 45,018 | 92,905 |
| hgnc | Human gene nomenclature and orthologies to mouse and rat | 42,741 | 38,360 |
| hgncgenefamily | Human gene-gene family memberships | 1,157 | 23,881 |
| hippie | Protein-protein physical interactions |  | 340,629 |
| homologene | Gene ortholog group memberships | 30,492 | 131,558 |
| hsdn | Disease-symptom associations |  | 10,246 |
| interpro | Protein-family and protein-domain memberships | 36,524 | 34,611 |
| kegg | Protein-pathway memberships | 330 | 30,346 |
| mgi | Mouse genome nomenclature | 300,499 |  |
| mirbase | MicroRNA nomenclature | 38,589 |  |
| mirtarbase | miRNA-target interactions |  | 366,110 |
| msig | Gene-gene set memberships | 17,810 | 2,443,391 |
| pfam | Protein-protein family and protein family-clan memberships | 17,929 |  |
| phewascatalog | Gene-disease relationships |  | 364,667 |
| phosphosite | Post-translational modifications |  | 553,716 |
| reactome | Protein-pathway and chemical-pathway memberships | 23,621 | 137,768 |
| rgd | Rat gene nomenclature | 44,970 |  |
| sider | Drugs' side effects and indications |  | 339,742 |
| wikipathways | Protein-pathway memberships | 513 | 22,115 |

While there are thousands of high quality databases available, including a high percentage that do not fit into the schemata defined by Pathway Commons, Bio2RDF, or other meta-databases that are more appropriate for BEL, we have prioritized them as they become have become relevant for our specific use-cases, but also are open to suggestions via the issue tracker on https://github.com/bio2bel/bio2bel/issues. Below, we present four of these use cases.

### 3.1. Mapping Concepts Between Pathway Databases with ComPath

Pathway databases have become one of the most frequently used biological data sources in the interpretation of high-throughput *-omics* experiments. Connecting pathway knowledge across the hundreds of databases developed in recent decades would not only provide a more comprehensive overview of the underlying biology they represent, but would also enable performing identical analyses on different databases. However, integrative approaches which combine databases lack the equivalence mappings between similar concepts and qualifiers that are necessary to compare between analyses run using one or another database. There are several reasons that explain the lack of mappings between databases, such as the absence of a common pathway nomenclature, differences in databases’ scopes, and the lack of clear pathway boundary definitions. Furthermore, generating high quality mappings requires a significant amount of manual effort since curators must individually investigate each pair of pathways and assess whether the pair comprises related or similar pathways occurring in the same biological context.

Three Bio2BEL packages were implemented for major pathway databases (i.e., KEGG, Reactome, and WikiPathways) and extended with tools to support the first curation of mappings between their equivalent and hierarchically related pathways during the ComPath project (Domingo-Fernández *et al.*, 2018). Each were used to store and harmonize the data underlying ComPath and its accompanying web curation interface (https://compath.scai.fraunhofer.de). Though the databases of the Bio2BEL packages are detached from the ComPath web application, they can be used to integrate additional biological data sources into ComPath in the future and also to regularly update their content over time (Wadi *et al.*, 2016); thus, facilitating the revisitation and reevaluation of the mappings.

### 3.2. Harmonizing Pathway Databases into a Common Schema with PathMe

The most direct and effective approach in addressing issues of interoperability of pathway databases is in the transformation of various database formats into a common schema. Although this approach has been exemplified by previously mentioned databases (e.g., OmniPath, Pathway Commons, and *graphite*; Sales *et al*., 2018), there have been several limitations which have impeded a complete harmonization of pathways from distinct biological data sources. Specifically, this requires: the harmonization of biological entities to identifiers from a common nomenclature (e.g., Entrez Gene or HGNC for human genes, ChEBI or PubChem for chemicals, etc.), the normalization of biological relationships, and an underlying format which serves as the unifying schema. However, a complete harmonization risks the loss of some information in the transformation process. For instance, pathway knowledge representations can span across several scales, such as molecular events, cellular processes, and phenotypes, which various formats accommodate for in varying degrees. While existing biological data sources can address certain aspects of these steps, addressing all of these steps would enable the complete interoperability of pathway databases. Accordingly, the PathMe software was designed to harmonize pathway databases into BEL as a common representation schema with Bio2BEL at its core (Domingo-Fernández *et al.*, 2019).

The selection of BEL lies in its flexibility to incorporate a wide range of biological entities from standardized nomenclatures and their relationships, all on a multi-modal scale. The transformation of various pathway formats into BEL through PathMe is facilitated by the Bio2BEL framework by allowing for the automation of the acquisition of the biological data sources which can change frequently. By integrating PathMe and Bio2BEL, any number of pathway resources included in the latter can be transformed into BEL. In doing so, users can enrich pathway knowledge by leveraging multiple, equivalent pathway representations from the various biological data sources included in Bio2BEL and analyze their own networks alongside canonical pathway ones. In a later publication, we plan to demonstrate the utility of combining Bio2BEL packages to produce an integrative pathway resource. Similarly to the recent comparison of pathway activity measurement tools by Lim *et al*. (2018), we will benchmark the performance of each of these resources both individually and combined on functional pathway enrichment and classification tasks applied to cancer genome and patient data.

### 3.3. Applications of Network Representation Learning with BioKEEN

The integration of numerous biological databases into a common schema gives rise to large, rich, heterogeneous knowledge graphs to which a variety of statistical and machine learning methodologies can be applied. One family of approaches, network representation learning (NRL), has been shown to be useful for clustering, entity disambiguation, and link prediction tasks (Nickel *et al.*, 2016). As new machine learning models are published for accomplishing these tasks, several implementations using the currently popular machine learning frameworks TensorFlow (Abadi *et al.* 2016) and PyTorch (Paszke *et al.*, 2017) provide reference implementations.

We developed BioKEEN as an extension to the previously developed NRL package, PyKEEN, to enable it to directly acquire and preprocess BEL knowledge graphs, namely those generated by Bio2BEL (Ali *et al*., 2018). One of the original goals of PyKEEN was to democratize NRL methods by facilitating those less familiar with the relevant mathematics and programming backgrounds to apply and evaluate them. We have continued this philosophy with BioKEEN to allow scientists to specify the Bio2BEL packages they would like to include in their analysis that are either hosted on PyPI, GitHub, or already installed as custom local packages. The usage of Bio2BEL allows scientists using NRL as a component of a more complex analytical pipeline to have the ability to not only re-run analyses in a reproducible manner, but also make use of the ability to acquire updated data when it becomes available.

Along with our previous publication, we provided several demonstrations including the prediction of novel protein-protein interactions using a model trained with the BioKEEN package for the Human Integrated Protein-Protein Interaction rEference (HIPPIE; Alanis-Lobato *et al.*, 2017), the prediction of pathway mappings using ComPath, and the prediction of disease-symptom associations using the Bio2BEL package for the HSDN (Zhou *et al.*, 2014) provided by Himmelstein *et al.* (2017) with Rephetio (https://het.io). Later, we plan to apply BioKEEN to combinations of Bio2BEL repositories to support other biologically relevant link prediction tasks such as drug repositioning.

### 3.4. Interoperability with Other Projects

The Integrated Network and Dynamical Reasoning Assembler (INDRA; Gyori *et al.*, 2017) integrates several databases including those covering physical interactions (e.g., BioGrid; Chatr-Aryamontri *et al.*, 2017), signaling (e.g., SIGNOR; Perfetto *et al*. 2016), curated drug targets (e.g., HMS LINCS small molecule target relationship database; http://lincs.hms.harvard.edu), and experimental drug affinities (e.g., Target Affinity Spectrum; Moret *et al*., 2018) in order to support generation of dynamical models. Following the recent development of a converter between BEL and INDRA (Hoyt *et al.*, 2019), these biological data sources can be indirectly made available as BEL, and all Bio2BEL packages can be integrated in INDRA.

Similarly, we are collaborating with the researchers developing OmniPath to structure their data acquisition pipelines as a Bio2BEL package, which is currently under development. Notably, OmniPath encompasses several biological data sources related to protein-protein interactions, transcriptional regulation, post-translational modifications, ligand-receptor interactions, and protein complexes, and others. This resource is complementary to content already available through Bio2BEL, providing a more comprehensive integration of the extensive publicly available biological data sources.

## 4. Conclusions

While the development of Bio2BEL has addressed the lack of defined schemata, data standardization, annotation of entities with classes, and application of controlled vocabularies to relations in numerous biological databases by converting them to BEL, several considerations remain. The approaches taken by Bio2RDF, Pathway Commons, and now Bio2BEL can be categorized as *data warehousing*. An alternative strategy, *data federation*, attempts to combine disparate biological data sources using SPARQL endpoints (e.g., DisGeNet-RDF (Queralt-Rosinach *et al.*, 2016), UniProt (Redaschi *et al.*, 2009), EBI (Jupp *et al.*, 2014)), RESTful APIs (e.g., BioServices (Cokelaer *et al.*, 2013), BioThings, Orange Bioinformatics (Curk *et al.*, 2005)), and more recently, GraphQL (https://graphql.org). Bio2BEL does not directly address data federation, but other aspects of the BEL ecosystem such as BEL Commons (Hoyt *et al.*, 2018b) have exposed RESTful APIs for manipulating BEL that might also be useful for GraphQL. However, the several attempts^12,13,14^ at converting BEL to RDF have suffered from relatively low adoption; and while a conversion to RDF enables querying with SPARQL, BEL lacks a dedicated query language that can leverage the rich aspects of its statements beyond their subjects, predicates, and objects.

Finally, it remains that like any format, consumers of BEL must make their own transformations appropriate for their scientific applications. We are not discouraged by this fact, and believe that Bio2BEL is a step towards enabling more computational scientists easy access to a larger portion of the wealth of available structured biological knowledge resources.

## 5. Availability and requirements

**Project Name**: Bio2BEL

**Project Home Page**: https://github.com/bio2bel

**Operating System(s)**: Platform independent

**Programming Language**: Python 3

**License**: MIT License

## Declarations

### Ethics Approval and Consent to Participate

Not applicable

### Consent for Publication

Not applicable

### Availability of Data and Materials

Each Bio2BEL package is listed on https://github.com/bio2bel and automatically acquires relevant data from their respective original biological data sources.

### Authors’ Contributions

CTH conceived and designed the study. CTH, DDF, and SM drafted the manuscript. MHA acquired funding and reviewed the manuscript. All authors performed data curation and developed computational pipelines for extraction, transformation, and loading of various biological data sources. All authors have read and approved the final manuscript.

## Acknowledgements

We would like to thank the curators and maintainers of the several databases we have used, without whom none of this work would be possible.

## Footnotes

1 https://www.w3.org/RDF

2 https://www.w3.org/TR/rdf-sparql-query

3 https://neo4j.com

4 https://orientdb.com

5 http://obofoundry.org/ontology/ro

6 https://psicquic.github.io/PSIMITAB.html

7 https://www.kegg.jp/kegg/xml/

8 http://sonnhammer.sbc.su.se/Stockholm.html

9 https://owlcollab.github.io/oboformat/doc/GO.format.obo-1_4.html

10 https://www.w3.org/OWL/

11 http://psidev.info/mif

12 https://wiki.openbel.org/display/OBP/BEL+RDF+Model

13 https://github.com/OpenBEL/bel2rdf

14 https://github.com/cthoyt/cx-rdf

## References

1. Abadi, M., et al. (2016). TensorFlow: A System for Large-scale Machine Learning. In Proceedings of the 12th USENIX Conference on Operating Systems Design and Implementation (pp. 265–283). Berkeley, CA, USA: USENIX Association. Retrieved from http://dl.acm.org/citation.cfm?id=3026877.3026899

2. Alanis-Lobato, G., Andrade-Navarro, M. A., & Schaefer, M. H. (2017). HIPPIE v2.0: enhancing meaningfulness and reliability of protein–protein interaction networks. Nucleic Acids Research, 45(D1), D408–D414. https://doi.org/10.1093/nar/gkw985

3. Ali, M., et al. (2018). BioKEEN: A library for learning and evaluating biological knowledge graph embeddings, 1–5. https://doi.org/10.1101/475202

4. Alocci, D., et al. (2015). Property Graph vs RDF triple store: A comparison on glycan substructure search. PLoS ONE, 10(12), 1–17. https://doi.org/10.1371/journal.pone.0144578

5. Belleau, F., Nolin, M.-A., Tourigny, N., Rigault, P., & Morissette, J. (2008). Bio2RDF: Towards a mashup to build bioinformatics knowledge systems. Journal of Biomedical Informatics, 41(5), 706–716. https://doi.org/10.1016/j.jbi.2008.03.004

6. Carbon, S., et al. (2017). Expansion of the gene ontology knowledgebase and resources: The gene ontology consortium. Nucleic Acids Research, 45(D1), D331–D338. https://doi.org/10.1093/nar/gkw1108

7. Cerami, E. G., et al. (2011). Pathway commons, a web resource for biological pathway data. Nucleic acids research. 39(Suppl. 1), D685–D690. https://doi.org/10.1093/nar/gkq1039.

8. Chatr-Aryamontri, A., et al. (2017). The BioGRID interaction database: 2017 update. Nucleic Acids Research, 45(D1), D369–D379. https://doi.org/10.1093/nar/gkw1102

9. Chen, B., et al. (2010). Chem2Bio2RDF: a semantic framework for linking and data mining chemogenomic and systems chemical biology data. BMC Bioinformatics, 11, 255.

10. Cokelaer, T., Pultz, D., Harder, L. M., Serra-Musach, J., & Saez-Rodriguez, J. (2013). BioServices: a common Python package to access biological Web Services programmatically. Bioinformatics, 29(24), 3241–3242. https://doi.org/10.1093/bioinformatics/btt547

11. Cote, R., et al. (2006). The Ontology Lookup Service, a lightweight cross-platform tool for controlled vocabulary queries. BMC Bioinformatics, 7, 1–7. https://doi.org/10.1186/1471-2105-7-97

12. Courtot, M., et al. (2011). Controlled vocabularies and semantics in systems biology. Molecular Systems Biology, 7(543). https://doi.org/10.1038/msb.2011.77

13. Curk, T., et al. (2005). Microarray data mining with visual programming. Bioinformatics, 21(3), 396–398. https://doi.org/10.1093/bioinformatics/bth474

14. Davidson, S. B., Overton, C., & Buneman, P. (1995). Challenges in integrating biological data sources. Journal of Computational Biology : A Journal of Computational Molecular Cell Biology, 2(4), 557–572. https://doi.org/10.1089/cmb.1995.2.557

15. Demir, E., et al. (2010). The BioPAX community standard for pathway data sharing. Nature Biotechnology, 28(12), 1308–1308. https://doi.org/10.1038/nbt1210-1308c

16. Domingo-Fernández, D., et al. (2018). ComPath: An ecosystem for exploring, analyzing, and curating mappings across pathway databases. npj Systems Biology and Applications, 4(1):43. https://doi.org/10.1038/s41540-018-0078-8.

17. Domingo-Fernández, D., Mubeen, S., Marin-Llao, J., Hoyt, C. T., & Hofmann-Apitius, M. (2019). PathMe: Merging and exploring mechanistic pathway knowledge. BMC Bioinformatics. In Press https://doi.org/10.1186/s12859-019-2863-9

18. Emon, M. A. E. K., Kodamullil, A. T., Karki, R., Younesi, E., & Hofmann-Apitius, M. (2017). Using Drugs as Molecular Probes: A Computational Chemical Biology Approach in Neurodegenerative Diseases. Journal of Alzheimer’s Disease, 56(2), 677–686. https://doi.org/10.3233/JAD-160222

19. Fabregat, A., et al. (2018). The Reactome Pathway Knowledgebase. Nucleic Acids Research, 46(D1), D649–D655. https://doi.org/10.1093/nar/gkx1132

20. Fan, X.-N., Zhang, S.-W., Zhang, S.-Y., Zhu, K., & Lu, S. (2019). Prediction of lncRNA-disease associations by integrating diverse heterogeneous information sources with RWR algorithm and positive pointwise mutual information. BMC Bioinformatics, 20(1), 87. https://doi.org/10.1186/s12859-019-2675-y

21. Gautier, L., Cope, L., Bolstad, B. M., & Irizarry, R. A. (2004). Affy - Analysis of Affymetrix GeneChip data at the probe level. Bioinformatics, 20(3), 307–315. https://doi.org/10.1093/bioinformatics/btg405

22. Gyori, B. M., et al. (2017). From word models to executable models of signaling networks using automated assembly. Molecular Systems Biology, 13(11), 954. https://doi.org/10.15252/msb.20177651

23. Himmelstein, D. S., Lizee, A., Hessler, C., Brueggeman, L., Chen, S. L., Hadley, D., … Baranzini, S. E. (2017). Systematic integration of biomedical knowledge prioritizes drugs for repurposing. ELife, 6. https://doi.org/10.7554/eLife.26726

24. Hoyt, C. T., et al. (2019). Re-curation and Rational Enrichment of Knowledge Graphs in Biological Expression Language. Database : The Journal of Biological Databases and Curation, baz068. https://doi.org/10.1093/database/baz068

25. Hoyt, C. T., Konotopez, A., & Ebeling, C. (2018a). PyBEL: a computational framework for Biological Expression Language. Bioinformatics, 34(4), 703–704. https://doi.org/10.1093/bioinformatics/btx660

26. Hoyt, C. T., Domingo-Fernández, D., & Hofmann-Apitius, M. (2018b). BEL Commons: an environment for exploration and analysis of networks encoded in Biological Expression Language. Database : The Journal of Biological Databases and Curation, 2018(3), 1–11. https://doi.org/10.1093/database/bay126

27. Irin, A. K., Tom Kodamullil, A., Gündel, M., & Hofmann-Apitius, M. (2015). Computational Modelling Approaches on Epigenetic Factors in Neurodegenerative and Autoimmune Diseases and Their Mechanistic Analysis. Journal of Immunology Research, 2015, 1–10. https://doi.org/10.1155/2015/737168

28. Iyappan, A., et al. (2014). NeuroRDF : Semantic Data Integration Strategies for Modeling Neurodegenerative Diseases. Proceedings of the 6th International Symposium on Semantic Mining in Biomedicine (SMBM2014), (January 2016), 11–18.

29. Iyappan, A., et al. (2017). Neuroimaging Feature Terminology: A Controlled Terminology for the Annotation of Brain Imaging Features. Journal of Alzheimer’s Disease, 59(4), 1153–1169. https://doi.org/10.3233/jad-161148

30. Jupp, S., et al. (2014). The EBI RDF platform: Linked open data for the life sciences. Bioinformatics, 30(9), 1338–1339. https://doi.org/10.1093/bioinformatics/btt765

31. Kanehisa, M., Furumichi, M., Tanabe, M., Sato, Y., & Morishima, K. (2017). KEGG: New perspectives on genomes, pathways, diseases and drugs. Nucleic Acids Research, 45(D1), D353–D361. https://doi.org/10.1093/nar/gkw1092

32. Laibe, C., & Le Novère, N. (2007). MIRIAM Resources: tools to generate and resolve robust cross-references in Systems Biology. BMC Systems Biology, 1, 58. https://doi.org/10.1186/1752-0509-1-58

33. Liberzon, A., et al. (2015). The Molecular Signatures Database Hallmark Gene Set Collection. Cell Systems, 1(6), 417–425. https://doi.org/10.1016/J.CELS.2015.12.004

34. Lim, S., Lee, S., Jung, I., Rhee, S., & Kim, S. (2018). Comprehensive and critical evaluation of individualized pathway activity measurement tools on pan-cancer data. Briefings in bioinformatics. https://doi.org/10.1093/bib/bby097

35. McKinney, W. (2010). Data Structures for Statistical Computing in Python. In S. van der Walt & J. Millman (Eds.), Proceedings of the 9th Python in Science Conference (pp. 51–56).

36. Meldal, B. H. M., et al. (2015). The complex portal - An encyclopaedia of macromolecular complexes. Nucleic Acids Research, 43(D1), D479–D484. https://doi.org/10.1093/nar/gku975

37. Menche, J., et al. (2015). Disease networks. Uncovering disease-disease relationships through the incomplete interactome. Science (New York, N.Y.), 347(6224), 1257601. https://doi.org/10.1126/science.1257601

38. Moret, N., et al. (2018). Cheminformatics tools for analyzing and designing optimized small molecule libraries. BioRxiv, (617), 358978. https://doi.org/10.1101/358978

39. Naz, M., Kodamullil, A. T., & Hofmann-Apitius, M. (2016). Reasoning over genetic variance information in cause-and-effect models of neurodegenerative diseases. Briefings in Bioinformatics, 17(3), 505–16. https://doi.org/10.1093/bib/bbv063

40. Nickel, M., Murphy, K., Tresp, V., & Gabrilovich, E. (2016). A Review of Relational Machine Learning for Knowledge Graphs. Proceedings of the IEEE, 104(1), 11–33. https://doi.org/10.1109/JPROC.2015.2483592

41. Paszke, A., Chanan, G., Lin, Z., Gross, S., Yang, E., Antiga, L., & Devito, Z. (2017). Automatic differentiation in PyTorch. 31st Conference on Neural Information Processing Systems, (Nips), 1–4. https://doi.org/10.1017/CBO9781107707221.009

42. Perfetto, L., et al. (2016). SIGNOR: A database of causal relationships between biological entities. Nucleic Acids Research, 44(D1), D548–D554. https://doi.org/10.1093/nar/gkv1048

43. Queralt-Rosinach, N., Piñero, J., Bravo, À., Sanz, F., & Furlong, L. I. (2016). DisGeNET-RDF: harnessing the innovative power of the Semantic Web to explore the genetic basis of diseases. Bioinformatics, 32(14), 2236–2238. https://doi.org/10.1093/bioinformatics/btw214

44. Redaschi, N., & Consortium, U. (2009). UniProt in RDF: Tackling Data Integration and Distributed Annotation with the Semantic Web. Nature Precedings. https://doi.org/10.1038/npre.2009.3193.1

45. Rogers, F. B. (1963). Medical subject headings. Bulletin of the Medical Library Association, 51, 114–6. Retrieved from http://www.ncbi.nlm.nih.gov/pubmed/13982385

46. Sales, G., et al. (2018). *meta*Graphite - a new layer of pathway annotation to get metabolite networks, Bioinformatics, bty719. https://doi.org/10.1093/bioinformatics/bty719.

47. Saqi, M., Lysenko, A., Guo, Y.-K., Tsunoda, T., & Auffray, C. (2018). Navigating the disease landscape: knowledge representations for contextualizing molecular signatures. Briefings in Bioinformatics, (May), 1–15. https://doi.org/10.1093/bib/bby025

48. Schriml, L. M., et al. (2018). Human Disease Ontology 2018 update: classification, content and workflow expansion. Nucleic Acids Research, 1–8. https://doi.org/10.1093/nar/gky1032

49. Slater, T. (2014). Recent advances in modeling languages for pathway maps and computable biological networks. Drug Discovery Today, 19(2), 193–198. https://doi.org/10.1016/j.drudis.2013.12.011

50. Stelzer, G., et al. (2016). The GeneCards suite: From gene data mining to disease genome sequence analyses. Current Protocols in Bioinformatics, 2016(June), 1.30.1-1.30.33. https://doi.org/10.1002/cpbi.5

51. Sun, K., Pržulj, N., Buchan, N., & Larminie, C. (2014). The integrated disease network. Integrative Biology, 6(11), 1069–1079. https://doi.org/10.1039/c4ib00122b

52. Szklarczyk, D., et al. (2015). STRING v10: Protein-protein interaction networks, integrated over the tree of life. Nucleic Acids Research, 43(D1), D447–D452. https://doi.org/10.1093/nar/gku1003

53. Türei, D., Korcsmáros, T., & Saez-Rodriguez, J. (2016). OmniPath: guidelines and gateway for literature-curated signaling pathway resources. Nature methods, 13(12):966. https://doi.org/10.1038/nmeth.4077.

54. van Dam, J. C. J., Schaap, P. J., Martins dos Santos, V. A. P., & Suárez-Diez, M. (2014). Integration of heterogeneous molecular networks to unravel gene-regulation in Mycobacterium tuberculosis. BMC Systems Biology, 8(1), 111. https://doi.org/10.1186/s12918-014-0111-5

55. Wadi, L., et al. (2016). Impact of outdated gene annotations on pathway enrichment analysis. Nature methods, 13(9):705. https://doi.org/10.1038/nmeth.3963.

56. Wanichthanarak, K., Fahrmann, J. F., & Grapov, D. (2015). Genomic, proteomic, and metabolomic data integration strategies. Biomarker Insights, 10 (Table 1), 1–6. https://doi.org/10.4137/BMI.S29511

57. Ward, L. D., & Kellis, M. (2012). HaploReg: A resource for exploring chromatin states, conservation, and regulatory motif alterations within sets of genetically linked variants. Nucleic Acids Research, 40(D1), 930–934. https://doi.org/10.1093/nar/gkr917

58. Warde-Farley, D., et al. (2010). The GeneMANIA prediction server: Biological network integration for gene prioritization and predicting gene function. Nucleic Acids Research, 38(SUPPL. 2), 214–220. https://doi.org/10.1093/nar/gkq537

59. Williams, A. J., et al. (2012). Open PHACTS: semantic interoperability for drug discovery. Drug Discovery Today, 17(21–22), 1188–1198. https://doi.org/10.1016/j.drudis.2012.05.016

60. Wishart, D. S., et al. (2018). DrugBank 5.0: a major update to the DrugBank database for 2018. Nucleic Acids Research, 46(D1), D1074–D1082. https://doi.org/10.1093/nar/gkx1037

61. Xin, J., et al. (2016). High-performance web services for querying gene and variant annotation. Genome Biology, 17(1), 91. https://doi.org/10.1186/s13059-016-0953-9

62. Yates, B., et al. (2017). Genenames.org: The HGNC and VGNC resources in 2017. Nucleic Acids Research, 45(D1), D619–D625. https://doi.org/10.1093/nar/gkw1033

63. Zhou, X., Menche, J., Barabási, A.-L., & Sharma, A. (2014). Human symptoms-disease network. Nature Communications, 5(May), 4212. https://doi.org/10.1038/ncomms5212

